# Semi-synthetic glycoconjugate vaccine candidate against *Cryptococcus neoformans*

**DOI:** 10.1101/2024.02.02.578725

**Authors:** Conor J. Crawford, Livia Liporagi-Lopes, Carolina Coelho, Samuel R. Santos Junior, André Moraes Nicola, Maggie P. Wear, Raghav Vij, Stefan Oscarson, Arturo Casadevall

**Affiliations:** Centre for Synthesis and Chemical Biology, University College Dublin, Belfield, Dublin, Ireland; Department of Molecular Microbiology and Immunology, Johns Hopkins Bloomberg School of Public Health 615 North Wolfe Street, Baltimore, MD 21205, USA; Max Planck Institute of Colloids and Interfaces, Am Mühlenberg1, 14476 Potsdam, Germany; Departamento de Análises Clínicas e Toxicológicas, Faculdade de Farmácia, Universidade Federal do Rio de Janeiro, Rio de Janeiro, Brazil; Faculty of Medicine, University of Brasília, Brasília, Brazil; MRC Centre for Medical Mycology, University of Exeter, Exeter Devon UK; Leibniz Institute for Natural Product Research and Infection Biology, Jena, Germany

## Abstract

*Cryptococcus neoformans* is a fungus classified by the World Health Organization as a critically important pathogen, posing a significant threat to immunocompromised individuals. In this study, we present the chemical synthesis and evaluation of two semi-synthetic vaccine candidates targeting the capsular polysaccharide glucuronoxylomannan (GXM) of *C. neoformans.* These semi-synthetic glycoconjugate vaccines contain the identical synthetic decasaccharide (M2 motif) antigen. This motif is present in serotype A strains, which constitute 95% of clinical cryptococcosis cases. This synthetic oligosaccharide was conjugated to two proteins (CRM197 and Anthrax 63 kDa PA) and tested for immunogenicity in mice. The conjugates elicited a specific antibody response that bound to the M2 motif but also exhibited additional cross-reactivity towards M1 and M4 GXM motifs. Both glycoconjugates produced antibodies that bound to GXM in ELISA assays and to live fungal cells. Mice immunized with the CRM197 glycoconjugate produced opsonic antibodies and displayed trends toward increased median survival relative to mice given a mock PBS injection (18 vs 15 days, *p* = 0.06). While these findings indicate promise, achieving a successful vaccine demands further optimization of the glycoconjugate. It could serve as a component in a multi-valent GXM motif vaccine, enhancing both strength and breadth of immune responses.

## Introduction

*Cryptococcus neoformans* is an environmentally ubiquitous fungus that can cause cryptococcosis in immunocompromised patients, such as solid-organ transplant recipients or individuals with AIDS.^1^ Infection often occurs undetected and the fungus can remain dormant in the host for decades in a state of latency.^1^ If immunosuppression occurs, new infections and latent infections can re-emerge, potentially leading to lethal meningitis. Current therapy necessitates the use of antifungal drugs for months, nonetheless resulting in high mortality, and clinicians are observing rising drug resistance.^2,3^ Hence, there is a need for better therapeutic and preventative strategies against cryptococcosis. The urgency of this issue was recognized by the World Health Organization (WHO), which listed *C. neoformans* in the critical priority group.^4^

Vaccination holds promise in preventing disease and eliminating the necessity for prolonged and costly antifungal treatments. In this regard, a prophylactic vaccine could potentially be used to safeguard at-risk groups, such as solid-organ transplant patients. A common strategy used in *C. neoformans* vaccine development has been to utilize antigens present on the fungal cell surface, including glycolipids, glycoproteins, and polysaccharides.^5–11^ The primary component of the *C. neoformans* capsular polysaccharide is glucuronoxylomannan (GXM), which is additionally essential for virulence.^12,13^ This polysaccharide does not contain a defined repeating unit but does contain discrete repeating motifs that occur together creating enormous antigenic and structural complexity (SI Figure 1).^14,15^ Importantly, to induce a protective immune response, a polysaccharide isolated from the microbial cell surface must be conjugated to a carrier protein, to provide T-cell help.^16^ Previous GXM-protein conjugate vaccines utilized biologically extracted native polysaccharides,^10,17^ which are highly heterogenous.^14^ Additionally, the conjugation method used, cyanogen bromide activation, is an imprecise chemical reaction that generates heterogeneous cross-linked matrices. *In vivo*, these conjugate vaccines were found to exhibit batch-to-batch variability, which could be attributed to the structural diversity of both the polysaccharides and the resulting conjugates. ^8,10^

In contrast, conjugate vaccines incorporating well-defined synthetic glycans offer distinct advantages. These include the ability to obtain a deeper understanding of how the mammalian immune system specifically recognizes carbohydrates, which, in turn, can allow rational design of vaccines. Potential benefits include enhanced immunogenicity, improved specificity, and more standardized composition. Synthetic carbohydrate antigens have been successfully used to develop commercial vaccines against *Haemophilus influenzae* type b and are under clinical investigation against *Shigella*.^18,19^ In addition, synthetic carbohydrate antigens serve as the antigenic component in candidate vaccines against fungi that have been tested in mice. This research has particularly concentrated on β-glucans,^20,21^ and β-mannans against Candida.^22,23^ A previous semi-synthetic glycoconjugate vaccine against *C. neoformans* utilized a heptasaccharide M2 motif antigen conjugated to human serum albumin (HSA) which induced an IgG immune response in mice.^11^ These antibodies recognized fungal cells in a punctuate pattern;^11^ however, monoclonal antibodies (mAbs) derived from the spleens of immunized mice were not opsonic towards fungal cells and did not protect mice in challenge experiments.^9^

Subsequent research, utilizing GXM microarrays,^24,25^ molecular modelling,^26^ and NMR spectroscopy,^27^ has revealed that larger M2 motif GXM oligosaccharides are required to adopt the conformations of GXM polysaccharides. Furthermore, these conformations are important for epitope presentation, enabling binding by both protective and non-protective mAbs.^24^ This means that the previously studied heptasaccharide M2 motif antigen was probably too small to assume solution-phase confirmations of GXM polysaccharides, potentially accounting for its poor efficacy *in vivo* and explaining the lack of binding to this structure on microarray surfaces. In contrast, decasaccharide **15** (Figure 1) has been shown to mimic the M2 motif in GXM polysaccharides and is widely recognized by protective and non-protective IgG and IgM mAbs on microarrays.^27^ Therefore, we hypothesized that **15** could be a more effective oligosaccharide antigen in vaccination.^25–27^ To test this hypothesis, we synthesized and evaluated two semi-synthetic vaccine candidates, both featuring the larger decasaccharide M2 motif conjugated to two distinct carrier proteins: CRM197 and anthrax protective antigen fragment 63 kDa (PA63) and evaluated their immunogenic properties in mice.

**Figure 1.**
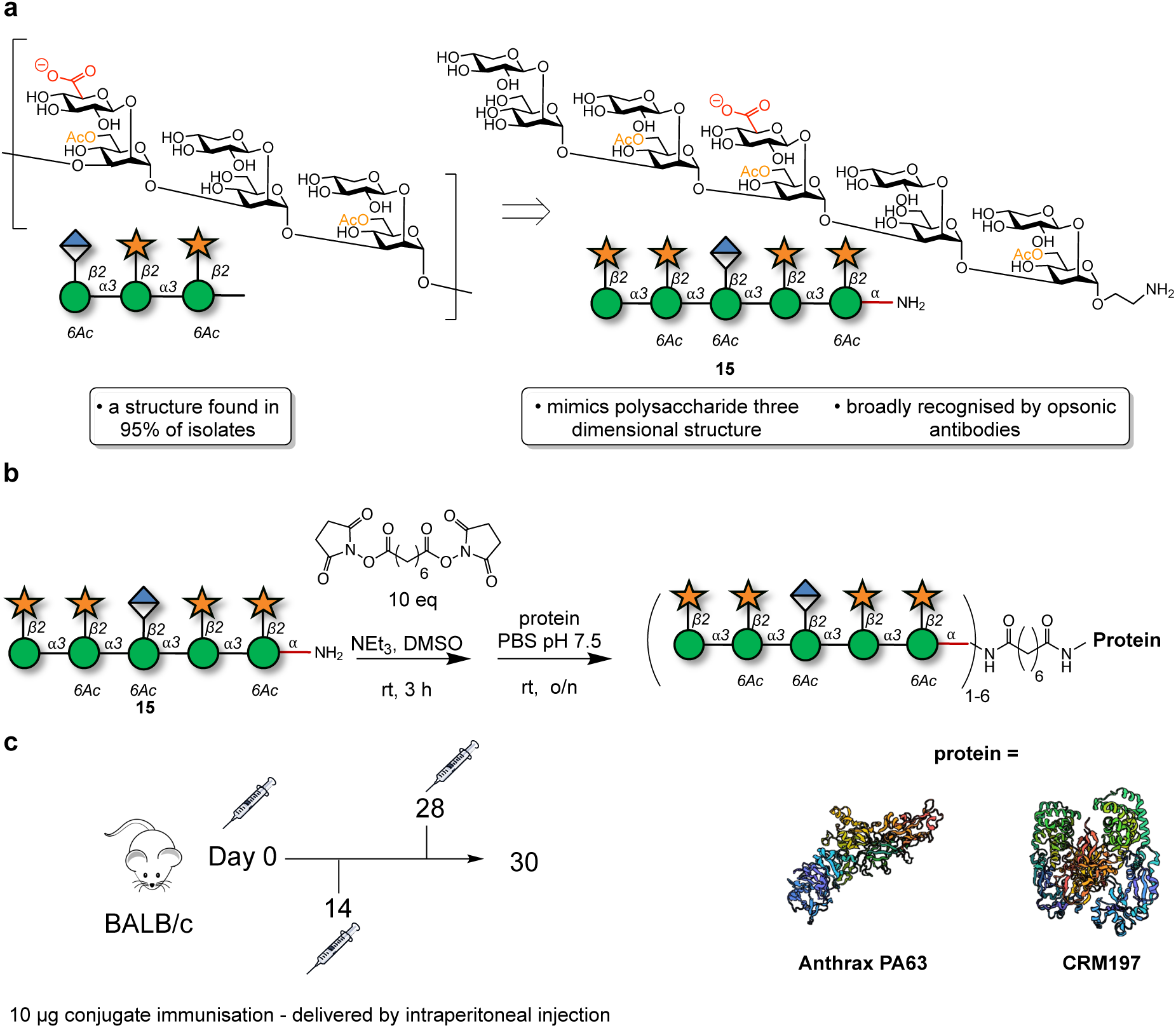
Synthesis and evaluation of two semi-synthetic vaccine candidates targeting the capsular polysaccharide, glucuronoxylomannan (GXM), of *C. neoformans*. **a.** Structure of motif 2 of GXM and the decasaccharide antigen. **b.** Conjugation of glycan **15** to carrier proteins anthrax PA63 or CRM197. **c.** Immunization schedule.

## Results and discussion

### Synthesis of semi-synthetic glycoconjugate vaccine candidates

The synthetic decasaccharide **15** was synthesized via a convergent approach employing thioglycoside building blocks.^25,28–30^ Following synthesis, glycan **15** underwent a reaction with an excess of a bis-NHS-activated suberic acid spacer. The intermediate product was then purified and conjugated to either CRM197 or PA63. MALDI-TOF analysis verified complete conversion of each carrier protein into glycoconjugates, with a loading of 1-6 glycans per protein (Figure 1, Table 1, SI Figure 2).

**Table 1.** Loading of DECA-Protein Conjugates.

| Entry | Antigen | Glycan:Protein | Protein | MW Range | Average Antigen Loading |
| --- | --- | --- | --- | --- | --- |
| 1 | **15** | 30:1 | CRM197 | 60-69 kDa | 3.5 |
| 2 | **15** | 30:1 | PA63 | 66-76 kDa | 3.5 |

### Immunization with synthetic decasaccharide-protein conjugates elicits antibodies that bind GXM

Mice (n = 5 control groups/PBS and n = 10 immunized groups) were intraperitoneally immunized with 10 μg of DECA-PA63 or DECA-CRM197 in complete Freund’s adjuvant, followed by two interval boosts 14 days apart (with 10 μg of conjugate in incomplete Freund’s adjuvant on days 14 and 28) (Figure 1c).

Subsequently, the immune serum reactivity with ELISA plates coated with GXM exopolysaccharide (EPS) confirmed a GXM-binding antibody response in mice after the third immunization in the DECA-CRM197 and DECA-PA63 groups, while no such response was observed in the control groups (Figure 2). The DECA-CRM197 sera bound to *C. neoformans* extracellular polysaccharide (EPS) with a similar affinity and decay pattern as mAb 18B7 (IgG1) (Figure 2a). Consequently, this serum was further examined for binding to CPS-coated ELISA plates. In contrast to the EPS-ELISA plates, mAb 18B7 exhibited a higher affinity (SI Figure 3). These results collectively suggest potential differences in the ratio of epitopes, quantity, and/or accessibility in the CPS and EPS preparations (SI Figure 3). Furthermore, competition ELISA with serum from DECA-CRM197 mice vs mAbs 18B7 (IgG1), 13F1 (IgM) and 2D10 (IgM) found that the DECA-CRM197 serum reducing binding of all three mAbs, implicating competition for the same epitopes (SI Figure 4). Moreover, a competition ELISA with serum from DECA-CRM197 mice versus mAbs 18B7 (IgG1), 13F1 (IgM), and 2D10 (IgM) revealed that the DECA-CRM197 serum reduced the binding of all three mAbs, implicating competition for the same epitopes (SI Figure 4).

**Figure 2.**
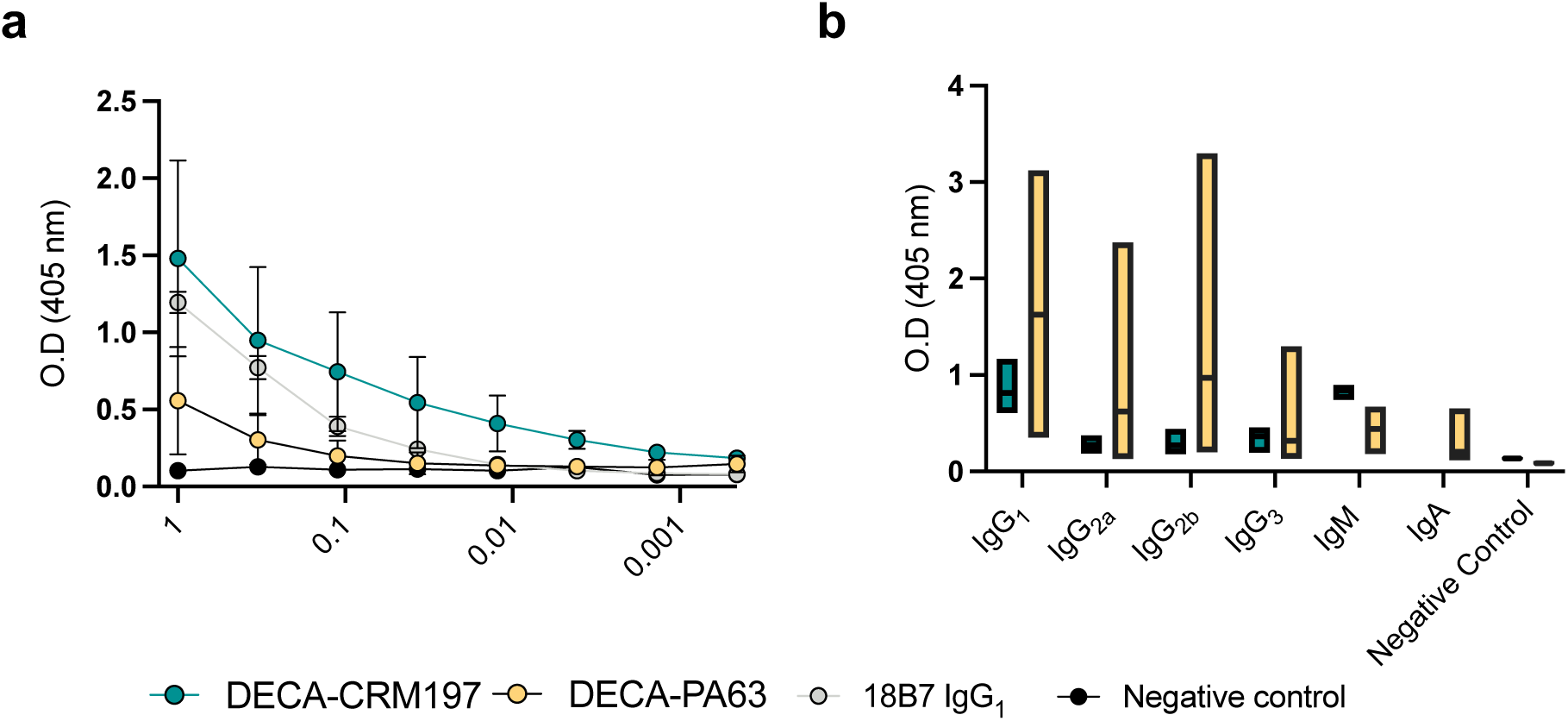
**a.** GXM ELISA of immunized mouse serum to EPS. For panel **a.** each dot represents the mean and bars represent standard deviation. A positive control of mAb 18B7 known to react with both GXM standards, and a negative control of PBS were used for the ELISA. **b.** Isotype distribution of antibodies found in sera from the DECA-CRM197 and DECA-PA63 conjugates. For isotype determination Indirect ELISA with CPS coating, followed by sera, and then antibodies of differing isotypes as indicated. All mice show a predominantly IgG1 and IgM isotypes. For panel **b.** the middle line in each bar indicates the mean, range is min to max. The serum for GXM reactivity and isotype analysis were isolated by retroorbital bleeding after the full immunization protocol. Experiments were repeated in triplicate.

### The isotype composition of antibodies in sera from glycoconjugate vaccine immunized mice was carrier protein-dependent

ELISA isotyping analysis of the immune sera from both conjugate vaccine immunizations uncovered a carrier protein dependent difference. While the overall immunoglobulin response being more consistent in DECA-CRM197 immunized mice compared to mice immunized with the DECA-PA63, while the later showed a greater range of affinity and isotypes (Figure 2b). The CRM197 immunized mice produced a predominance of IgG_1_ and IgM isotype classes with lower levels of IgG2a, IgG2b and IgG3 detected. In the DECA-PA63 immunized mice high levels IgG1 were also detected but an additional high level of the IgG2b subclass. The DECA-PA63 mice also produced lower levels of IgM, IgG2a and IgA. These results highlight the importance of the carrier.^31^

### Immunization with GXM M2 motif elicits cross-reactive sera

The molecular reactivity of the antibodies elicited during immunization was analyzed using a GXM microarray by selecting three mice randomly from each group.^24^ The microarray analysis revealed that the conjugates predominantly elicited an immune response specific to the M2 motif of *O*-acetylated glycans (**15**–**18**, **25**), with comparatively weak reactivity to other GXM motifs. A single mouse (n = 6) immunized with the DECA-PA63 conjugate elicited antibodies that manifested reactivity toward the non-*O*-acetylated decasaccharide **25** (Figure 3b, row A). No binding to M2 motif glycans smaller than the decasaccharide **15** were observed in the DECA-CRM197 mice. In contrast, the DECA-PA63 conjugate elicited antibodies with reactivity towards two smaller structures: the first, a disaccharide of glucuronic acid-β-(1,2)-linked to 6-*O*-acetylated mannose (**2**), a motif found in the center of the decasaccaride antigen; the second, tetrasaccharide **11**, the terminal epitope of the decasaccaride antigen (Figure 3a).

**Figure 3.**
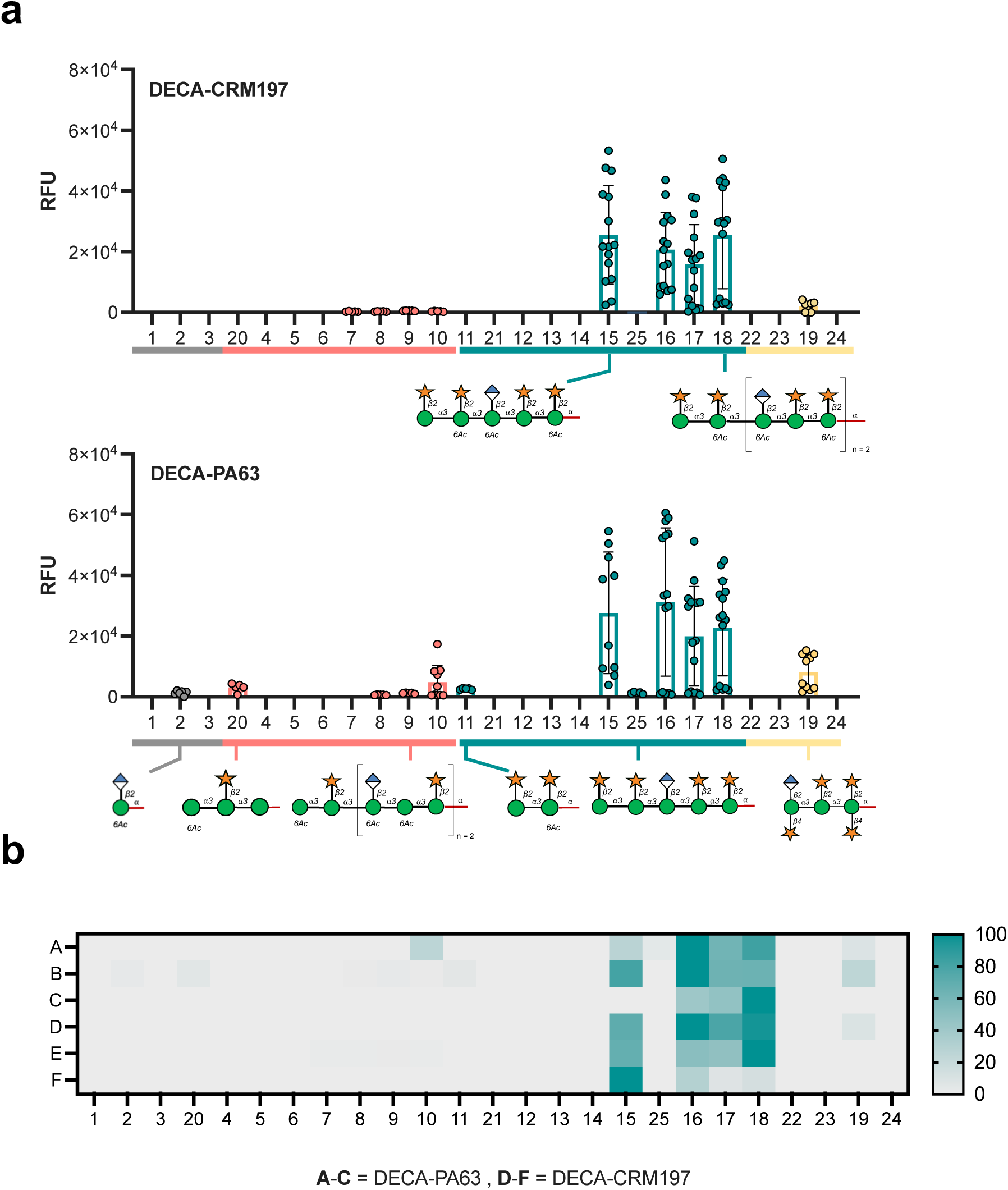
Microarray reactivity analysis of sera from mice immunized with CRM197 and PA63 glycoconjugates towards oligosaccharides. **a.** Pooled response of DECA-CRM197 and DECA-PA63 mice **b.** A heatmap of comparing glycan binding profiles of sera from both conjugates. Rows A-C are DECA-PA63 and rows D-F are DECA-CRM197 mice. The heatmap is normalized by lowest and highest point in each dataset. The full library of the oligosaccharides structure can be found in SI Figure 5. A data point is the RFU of a single glycan spot on the array.

Cross-reactivity toward other GXM motifs was observed for antibodies elicited by both GXM conjugates. Both conjugates elicited reactivity towards glycan **19**, a non-acetylated octasaccharide representative of the M4 motif. Additionally, the glycoconjugates elicited additional reactivity towards the M1 motif, for the DECA-PA63 mice recognizing structures **8**, **9**, **10**, and **20**. While, the DECA-CRM197 sera has reactivity towards M1GXM motif glycans **7**-**10** (Figure 3). Surprisingly, a single mouse from the DECA-PA63 group (total n of mice = 3) did not recognize decasaccharide **15** on the microarray but did show reactivity towards larger M2 motif glycans **16**, **17** and **18** (Figure 3b, row C). These results demonstrate heterogeneity in the specificity of the immune response, even among genetically identical mice.

The binding reactivities of mAbs 18B7, 13F1, and 2D10 were previously determined by GXM microarray, indicating a predominant affinity towards the M2 motif of GXM.^24^ Moreover, 18B7 and 13F1 displayed additional reactivity towards the M4 motif, and 18B7 demonstrated the ability to bind to the M1 motif. Therefore, the microarray analysis supports the findings of the competition ELISA (SI Figure 4), suggesting that the elicited antibodies could compete for M1, M2, and M4 motifs.

The low cross-reactivity observed in mice immunized with a single GXM motif conjugate suggests that all GXM motifs, when present in large oligosaccharides, share common structural epitopes. This is evident from the rare binding to GXM M2 oligosaccharides smaller than decasaccharide (**15**), and that cross-reactivity between motifs only initiates at the decasaccharide (**7**) level for M1 motifs and at the octasaccharide stage for GXM M4 motifs (Figure 5a). For future vaccine candidates, it would be desirable to elicit reactivity towards all GXM motifs at high levels. This implies the necessity for incorporating these oligosaccharides into conjugate vaccines.

### Semi-synthetic glycoconjugates elicit antibodies that bind to *C. neoformans* capsules

The capsule of *C. neoformans* displays remarkable antigenic diversity, characterized by a range of GXM motifs (SI Figure 1).^32^ These motifs exhibit dynamic variations both spatially, in their proximity to the cell wall, and temporally, evolving over the fungus’s lifecycle.^33^ The binding patterns of antibodies to the capsule have implications for the protective and non-protective efficacy of monoclonal antibodies.^34^ These binding patterns are intricately linked to the abundance and location of epitopes within the capsule, as well as the antibodies’ capacity to bind these epitopes.

Our investigation into antibodies elicited by semi-synthetic conjugates involved using sera obtained post a full immunization protocol for live imaging of fungal cells. For immunofluorescence, we utilized two strains of C. neoformans: H99, a serotype A lab strain known to express multiple GXM motifs, and Mu-1, also serotype A, which exclusively expresses the M2 motif of GXM.^14^ Notably, the serum binding to H99 fungal cells from the DECA-PA63 conjugate showed low consistency (8%, 4 positive cells, 49 cells total), with the majority of cells showing no staining. In contrast, the DECA-CRM197 conjugate elicited antibodies that stained the majority of cells (80%, 20 positive cells, 24 cells total) (Figure 4), potentially could reflecting its higher immunogenicity (Figure 4). Interestingly, both vaccine sera exhibited a punctate pattern when binding to H99, while binding to the Mu-1 strain displayed a more annular pattern. Previous analyses have shown that mAb 18B7 yields annular staining of H99, whereas 2D10 staining is punctate.^35^

**Figure 4.**
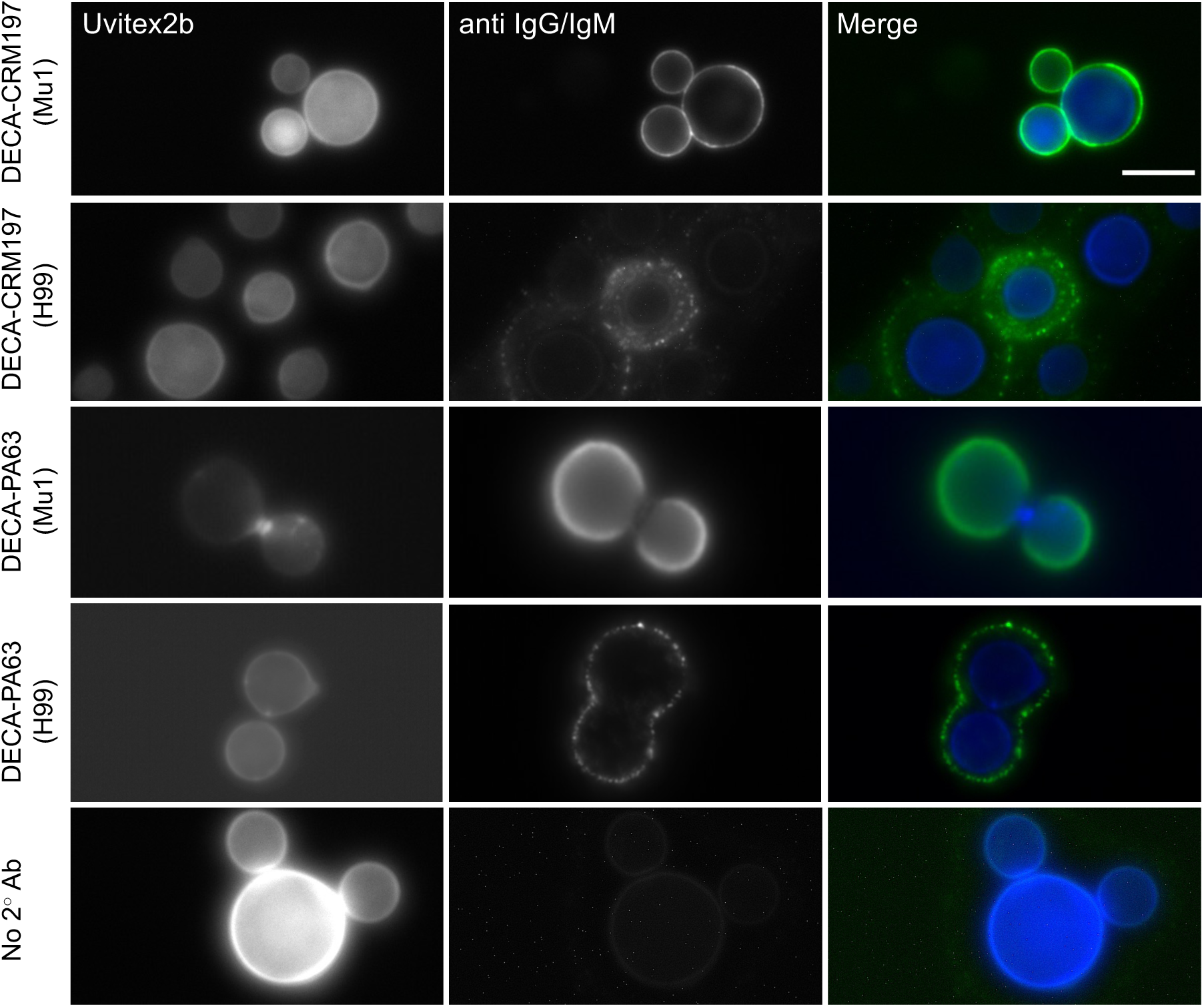
Antibodies from semi-synthetic conjugates bind to *C. neoformans* cells. Mu1 and H99 were cultured for 48 h in capsule-inducing media and immunofluorescence was performed using the mouse serum from mice treated with 10µg DECA-CRM197 or 10µg DECA-PA63 conjugate. Cells were imaged for Uvitext2b (Blue channel), secondary antibodies (IgG and IgM) (Green channel). Left margin indicates conjugate and inside the parentheses is the strain of *C. neoformans*. Merged image show annular and punctate staining in immunized mouse serum dependent on strain. Scale bar is 10 µM.

**Figure 5.**
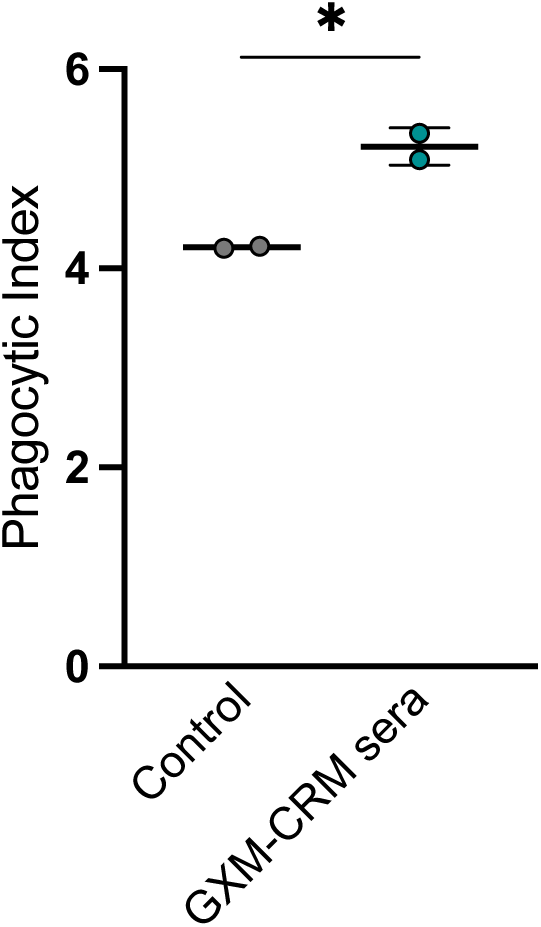
The DECA-CRM197 conjugate vaccine elicited opsonic antibodies. The dot represents an individual experiment, the line is the mean and error bars are standard deviation (SD). BMDM-derived macrophages were co-cultured with *C. neoformans* strain 24067 in the presence of 20% guinea pig serum (as a control) and 1:50 dilution of serum from 10 µg immunized mice. Cells were counted after 2 hours by using microscopy with a 40x objective. The phagocytic index (MOI 3:1) was determined by the number of internalized cryptococcal cells per 100 macrophages.

This observation is supported by microarray analysis, which demonstrates that conjugates primarily are reactive to the M2 motif (Figure 3). This suggests that achieving annular binding to antigenically diverse capsules requires antibodies with both diverse and high affinity binding, necessitating immunization with several GXM motifs (Figure 6).

**Figure 6.**
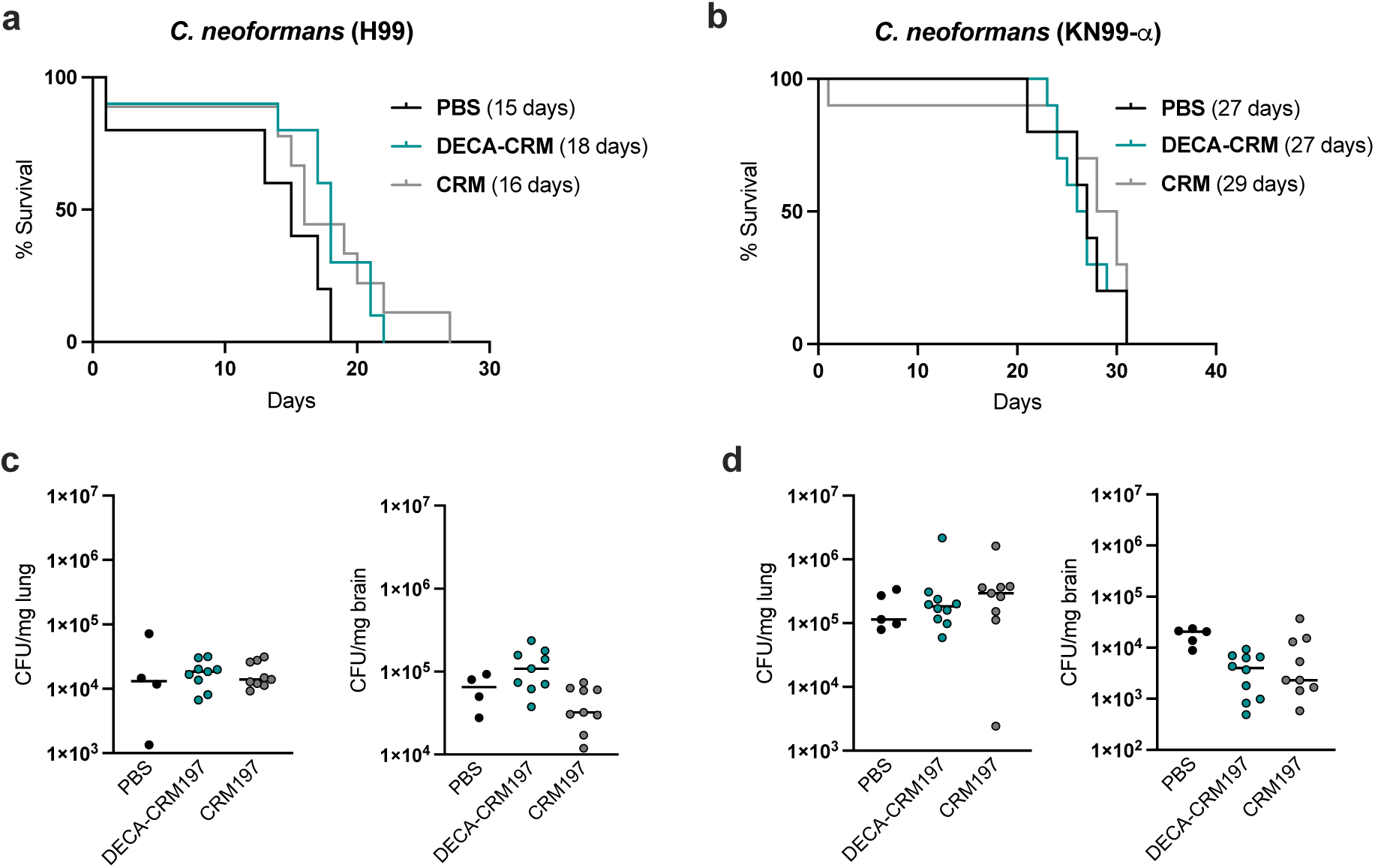
Challenge experiments with DECA-CRM197 conjugates against two strains of *C. neoformans*. **a.** H99 challenge and **b.** KN99-α challenge, in parentheses is the median survival time. **c.** Lung and brain fungal burden at day of the mouse death via CFU for H99 **d.** Lung and brain fungal burden at day of the mouse death via CFU for KN99-α challenge. Shown is median with individual points showing CFU per tissue/mouse.

### The DECA-CRM197 conjugate vaccine elicits opsonic antibodies

The capacity of antibodies to promote phagocytosis by immune cells is linked to protection in cryptococcosis.^36^ The DECA-CRM197 sera was subject to further investigation for its ability to phagocytose fungal cells because of : (i) its higher binding by ELISA (Figure 2a) and (ii) the microarray analysis suggested that these sera bound to conformational epitopes, which has been shown to be crucial for mAbs known to be opsonic (Figure 3a). (iii) and its more reliable binding to fungal cells by immunofluorescence (Figure 4). The sera derived from DECA-CRM197 immunization increased phagocytosis by BMDM cells (1:50 dilution) (*, *p* = 0,0169, unpaired t-test) against *C. neoformans* compared to the control group (20% (v/v) guinea pig serum) (Figure 5). This sets the DECA-CRM197 conjugate apart from previous heptasaccaride-HSA conjugate which did not illicit phagocytic antibodies^9,11^ This distinction may arise due to: (i) the different glycan antigens in the conjugates or (ii) the use of a more immunogenic carrier protein.

Overall, the increase in opsonic efficacy for vaccine immune sera relative to the control was relatively small, and it is important to note that opsonic antibodies are not necessarily protective. This is evidenced by the efficiency of a non-protective IgG3 to GXM as an opsonin.^37^

### Immunization DECA-CRM197 had modest or no effect on survival time

The immunization protocol was repeated (10 µg x 3, days 14, 21, 28) and on day 42 the mice (n = 5 control groups and n = 10 immunized groups) were challenged with 1.0 x 10^7^ yeasts of *C. neoformans* strain H99 or KN99-α.

Mice challenged with *C. neoformans* H99 had the longest median survival time when they received the DECA-CRM197 conjugate (18 days). Followed by those immunized with CRM197 (16 days), and those given a PBS mock infection had the lowest median survival time (15 days) (Figure 6a). Comparison of mice immunized with DECA-CRM197 to those given a mock PBS infection was just above significance (*p* = 0.0678, Kaplan-Meier survival analysis Log-rank (Mantel-Cox) test).

Post challenge all mice were analyzed for colony forming units (CFUs) in the lung and the brain, which are organs known to be involved in systemic cryptococcosis. Finding that CFUs in the brain of H99 challenged mice show a significant difference between groups (** p = 0,0052, Kruskal-Wallis test). CRM197 immunized mice were found to have lower CFUs compared to the DECA-CRM197 mice (**, p = 0.0078, post-hoc Dunn’s multiple comparisons test) (Figure 6c), raising the possibility that some of the effects observed in mice reflected a protective effect to the carrier protein. Analysis of the colony forming units (CFUs) in the lungs showed no difference between groups (ns, p = 0.8217, Kruskal-Wallis test).

Mice challenged with strain KN99-α showed no difference in survival (n = 5 control groups and n = 10 immunized groups, Kaplan-Meier survival analysis), with CRM immunized mice having the highest median survival time at 29 days, and mice in both the DECA-CRM197 and PBS having median survival times of 27 days (Figure 6b).

In the KN99-α challenged mice, the groups showed distinct response in the brain (*, p = 0.0193, Kruskal-Wallis test) with the DECA-CRM197 immunized mice having lower CFUs compared to control groups (*, p = 0.0172, post-Dunn’s test), while in the lungs, KN99-α challenged mice showed no distinct response between groups (ns, p = 0,3898, Kruskal-Wallis test) (Figure 6d).

## Conclusions

We report the synthesis and evaluation of two semi-synthetic vaccine candidates targeting *C. neoformans*, both incorporating a decasaccharide (DECA) antigen but utilizing different carrier proteins (CRM197 and PA63). Mice exhibited a carrier protein-dependent immune response, with the DECA-CRM197 conjugate demonstrating a more defined and therefore more consistent outcome in mice. The DECA-CRM197 conjugate induced primarily IgG1 and IgM antibodies, while the DECA-PA63 conjugate induced IgG1 and IgG2b antibodies. Glycan microarray analysis revealed that both conjugates predominantly elicited sera reactivity towards the M2 motif, with concurrent weak cross-reactivity with the M1 and M4 motif. Sera from immunized mice had the ability to bind to *C. neoformans* H99 cells in a punctate pattern, which is associated with non-protective antibodies in murine challenge studies, but did bind in annular patterns to the M2 motif capsule of Mu-1 cells.^34^ A key distinction over the previous heptasaccharide semi-synthetic vaccine candidate was that the DECA-CRM197 immunized mice sera was opsonic although the opsonic efficacy of the immune serum was modest.

Despite promising aspects, the challenge experiment revealed only a modest increase in median survival time against strain H99, with no statistical significance against KN99-α. Potential factors contributing to this outcome include inadequate levels of protective antibodies, low protective efficacy of the antibody response, and uncertainties around optimal glycan-protein loading, formulation, and immunization schedules. Furthermore, the punctate binding of mice sera to H99 cells observed through immunofluorescence implies that these cells would likely escape the immune response, possibly accounting for the modest protection observed. Overall, the DECA-CRM197 conjugate demonstrated enhanced efficacy compared to a previous heptasaccharide vaccine,^11^ but the results still fall short of those reported with glycoconjugates using cryptococcal GXM.^38^ The antigenic diversity of the GXM polysaccharide suggests a need for a multivalent approach, encompassing each motif of GXM. Our findings highlight the potential of the DECA-CRM197 conjugate as a valuable component in future multivalent vaccines aimed at preventing cryptococcosis.

## Author Contributions

C.J.C, S.O and A.C wrote the original draft. C.J.C completed the synthesis and the conjugations. S.R., L.L.L, A.M.N, C.C and C.J.C completed animal immunizations. C.J.C printed and screened the glycan arrays. S.R, L.L.L carried out challenge experiments. C.J.C and R.V completed the immunofluorescence experiments. All authors designed and planned the study and edited manuscript. C.J.C, S.O and A.C funding acquisition.

## Acknowledgments

We thank Dr Yannick Ortin for NMR support. C.J.C was funded by Irish Research Council postgraduate award (GOIPG/2016/998). MPW was supported in part by AI007417. S.O was supported by Science Foundation Ireland Award 13/IA/1959 and 20/FFP-P/884. AC was supported in part by NIH grants AI052733-16, AI152078-01 and HL059842-19.

## Methods

### Oligosaccharide synthesis

Chemical synthesis of decasaccharide **15** followed published procedures.^25,28,29,39,40^ We used a convergent synthesis of di- and tetrasaccharide thioglycoside building blocks, which were coupled using dimethyl(methylthio)sulfonium trifluoromethanesulfonate (DMTST) in diethyl ether.^41^ A 2-naphthylmethyl (NAP) ether protecting group was employed as a temporary protecting group to allow extension of the oligosaccharide from the non-reducing end. Cleavage of the NAP ether was complete using 2,3-dichloro-5,6-dicyano-1,4-benzoquinone (DDQ) with a buffered aqueous component to suppress cleavage of benzyl ethers.^28^ Deprotection of the GXM used a pre-conditioned palladium on carbon catalyst to which gave a selective catalyst for hydrogenolysis of the aromatic protecting groups yield the desired decasaccharide **15**.^28,42^

### Conjugations

Decasaccharide **15** (1 eq) was coupled to a bis(*N*-hydroxysuccinimide ester) suberic acid linker (5 eq) in DMSO, which was followed by MALDI-TOF (SuperDHB, acetonitrille:water – 1% TFA, 50:50 v/v). Once complete, 5**1** was precipitated by addition of ethyl acetate and centrifuged at 4 °C for 5 min. The supernatant was removed and washed twice with cold ethyl acetate to remove excess suberic acid linker. The precipitate was then resuspended in PBS (100 mM, pH 7.5) and added in 30-fold excess to either a solution of CRM197 or PA 63 (PBS (100 mM, pH 7.5) and left overnight at room temperature (30 glycan: 1 protein). Conjugates were followed and analyzed by MALDI-TOF spectrometry in linear-mode to determine the extent of conjugation (*trans*-ferulic acid, ethanol: water - 0.5% TFA, 50:50 v/v). Once satisfactory loading was achieved, the reaction mixture was transferred to a Vivaspin® 500 centrifugal filter (MWCO 10 kDa, GE Healthcare, Buckinghamshire, UK), desalted and washed three times with sterile water (MIKRO 200R, Hettich®, 15 minutes at 13000 RPM, 3 x 500 μL water). The conjugate was suspended in 1 mL sterile water and final concentration of protein was determined using the extinction co-efficient (ε) ε0.1% 1.07 for a 1 mg/ml at 280 nm (UV-1280 Shimadzu spectrometer) to yield 2.0 mg of the conjugate. Finally, the conjugate was lyophilized to give a white solid.

### Ethics statement

All animal procedures were performed with prior approval from Johns Hopkins University (JHU) Animal Care and Use Committee (IACUC), under approved protocol number MO18H152. Mice were handled and euthanized with CO_2_ in an appropriate chamber followed by thoracotomy as a secondary means of death in accordance with guidelines on Euthanasia of the American Veterinary Medical Association. JHU is accredited by AAALAC International, in compliance with Animal Welfare Act regulations and Public Health Service (PHS) Policy and has a PHS Approved Animal Welfare Assurance with the NIH Office of Laboratory Animal Welfare. JHU Animal Welfare Assurance Number is D16-00173 (A3272-01). JHU utilizes the United States Government laws and policies for the utilization and care of vertebrate animals used in testing, research and training guidelines for appropriate animal use in a research and teaching setting.

### Microorganisms and growth conditions

*C. neoformans* serotype A strains H99 (ATCC 208821) and Kn99α were used in the DECA-CRM197 challenge experiments. The yeast cells were kept frozen in 10%-glycerol. Sabouraud dextrose broth (SAB, from Gibco) medium was used for standard growth of yeast cells at 30 °C with moderate shaking (120 rpm) overnight. For the immunofluorescence experiments, after the preculture on SAB medium, *C. neoformans* cells were grown for 3 days at 30 °C/120 rpm in minimal media (15 mM dextrose, 10 mM MgSO_4_, 29.4 mM KH_2_PO_4_, 13 mM glycine, and 3 μM thiamine-HCl), to maximize capsule development.

#### Cell culture

NSObcl2 cells were obtained from the Albert Einstein College of Medicine hybridoma facility and maintained in RPMI supplemented with 10% fetal calf serum and 1 µg/mL G418.^43^ Hybridomas generated as described below were cultivated in the same medium.

### Mice and immunization

Six-week-old female A/J mice were used to perform the immunizations. Mice were immunized intraperitoneally with 10 μg of DECA-PA63 or DECA-CRM197 in complete Freund’s adjuvant (Sigma, St. Louis, MO). Mice were boosted at 14 days interval or as required with 10 μg of each conjugate in incomplete Freund’s adjuvant.^9^ Mice were bled by retro-orbital bleeding using heparin capillary tubes under isoflurane anaesthesia, two weeks after immunizations.

### Serological assays

Immune sera were assayed for their reactivity against GXM by indirect Enzyme-Linked Immunosorbent Assay (ELISA). Exopolysaccharides (EPS) obtained by ultrafiltration from *C. neoformans* strain H99 cultures in minimal media grown for 3 days in a concentration of 10 μg/mL, were dissolved in PBS used to coat polystyrene plates (Corning *9018) by incubating overnight at 4°C. CPS isolation and ELISA plates were created following published protocols.^44^ After blocking with 1% BSA in PBS, dilutions of the sera from the immunized mice were incubated for 1 h at 37°C and then overnight at 4°C. As secondary antibody we used alkaline phosphatase-conjugated goat anti-mouse IgG, IgA, and IgM (Southern Biotech) in a 1:1000 dilution, as indicated in figure legends, for 1 h at 37°C. In other experiments, we used as secondary antibodies isotype-specific (IgA, IgG1, IgG2a, IgG2b, IgG3) alkaline-phosphatase-conjugated antibodies (Southern Biotech). Bound antibodies were detected using p-nitrophenyl phosphate (Sigma) as a substrate, with absorbance measured in a plate spectrophotometer at 405 nm. For competition ELISA assays mAbs 18B7, 13F1 and 2D10 were biotinylated with a biotin commercial kit, according to the manufacturer’s instructions (Pierce, Rockford, IL, USA). ELISA plates were generated as described above. After blocking, a constant concentration of the biotinylated mAb was incubated with decreasing concentrations of a different non-biotinylated mAb in blocking buffer for 1 h at 37°C. After washing, avidin conjugated with alkaline phosphatase (Sigma-Aldrich) was added, and the preparation was incubated for 1 h at 37°C. Absorbance at 405 nm was recorded after the reaction was developed with *p*NPP.^45^

### Glycan Microarray Scanning

The glycan microarray scanning was carried out as described.^46^ Primary anti-GXM mAbs or control Abs were prepared from stocks to the necessary concentration in 3% BSA in PBS-T. Biotinylated goat anti-mouse kappa chain Abs were used as secondary reagents for all primary antibodies. Detection was performed with the streptavidin-conjugated SureLight P3 fluorophore (Cayman Chemical Company, Ann Arbor, MI) at 5 μg/mL in PBS-T. All hybridization steps were performed using the Agilent 8-well gasket system in a humidity-controlled rotating hybridization oven at 26° C for 1-2 hours. Washes (X3) in Tris-buffered saline (pH 7.6, 0.1% Tween 20) (TBS-T) for 3 minutes and once for 3 minutes in TBS. Scanning was performed in an Agilent SureScan Dx microarray scanner with red wavelength emission detection. The data was processed on Mapix software. The mean fluorescent intensities (corrected for mean background) and standard deviations (SD) were calculated (n=6).

### Phagocytosis assay

Phagocytosis assay was performed as described: BMDM were seeded (5 × 10^4^ cells/well) on poly-D-lysine coated coverslip bottom MatTek petri dishes with 14 mm microwell (MatTek Brand Corporation). Cells were then incubated at 37°C with 10% CO_2_ overnight. On the following day, BMDMs were infected Uvitex 2B (Polysciences, Warrington, PA) stained with cryptococcal cells (1.5 × 10^5^ cells/well), and with addition of sera (1:50 (v/v)) from three different mice of the same immunization group and/or complement (20% (v/v)) guinea pig serum - Fisher Scientific #642831). After 2 h incubation to allow phagocytosis, culture was washed five times with fresh medium to remove extracellular cryptococcal cells. In addition, Alexa fluor 568 (Thermo Fisher Scientific) conjugated to 18B7 mAb was added to stain remaining extracellular fungal cells. Images were taken by using Olympus AX70 microscopy (Olympus, Center Valley, PA) with a 40x objective. The *C. neoformans*/macrophage ratio was 3:1. The phagocytic index was determined by the number of internalized cryptococcal cells per 100 macrophages.^47^

### Immunofluorescence

Cells were grown in minimal media for 3 days. The cells were then washed twice in PBS and counted using a hemocytometer. Approximately 2.5 x 10^5^ cells/ ml in 100 µL of blocking solution (PBS-1% BSA) was incubated with sera at a dilution of 1:50 for 30 minutes at RT. Cells were washed twice in blocking solution. 1:100 of secondary antibodies; Goat anti rat IgM (1 mg/ ml), Goat anti mouse kappa IgG FITC (1 mg/ ml), Rabbit anti goat IgG FITC (1 mg/ ml) and Goat anti mouse IgM (1 mg/ ml), and 0.1 µg/ mL Uvitex 2B, were added to the cells. Cells were imaged using Olympus AX70 microscope.

### Survival study

Mice (n = 7 animals in control groups and n = 8 animals treated groups) were immunized three times (day 0, day 14, and day 28) as follows: (1) a group was injected intraperitonially only with CRM197 (control group #1); (2) a group was injected intraperitonially with DECA-CRM197 conjugate, and (3) a group was injected intraperitonially with PBS (control group #2). All immunogens were emulsified in Freund’s adjuvant (first one with complete adjuvant; second and third ones with incomplete adjuvant). Mice were challenged intranasally with 1.0 × 10^7^ *C. neoformans* cells per animal (KN99- or H99)^48^ 14 days after last immunization. The animals were observed daily for 30 days and euthanized if showing more than 20% weight loss or inability to feed.

### Fungal burden assessment

The fungal burden was evaluated at the time of death by counting CFU (colony-forming units). Lungs and brain were removed, weighed and homogenized in 1 mL of PBS. After serial dilutions, homogenates were inoculated on Sabouraud agar plates with 1% streptomycin/penicillin (Corning, NY). The plates were incubated at room temperature, and the colonies were counted after 48-72 h.

### Statistics

Statistical analyses were done using GraphPad Prism version 8.00 for Mac OS X (GraphPad Software, San Diego, CA, USA). Statistical analyses for the survival analysis used Kaplan-Meier with Log-rank (Mantel-Cox test) and Gehan-Breslow-Wilcoxon tests. One-way analysis of variance using a Kruskal-Wallis nonparametric test was used to compare the differences between groups, and individual comparisons of groups were performed using a Dunn’s multiple comparisons test.

## Supporting Figures

**SI Figure 1.**
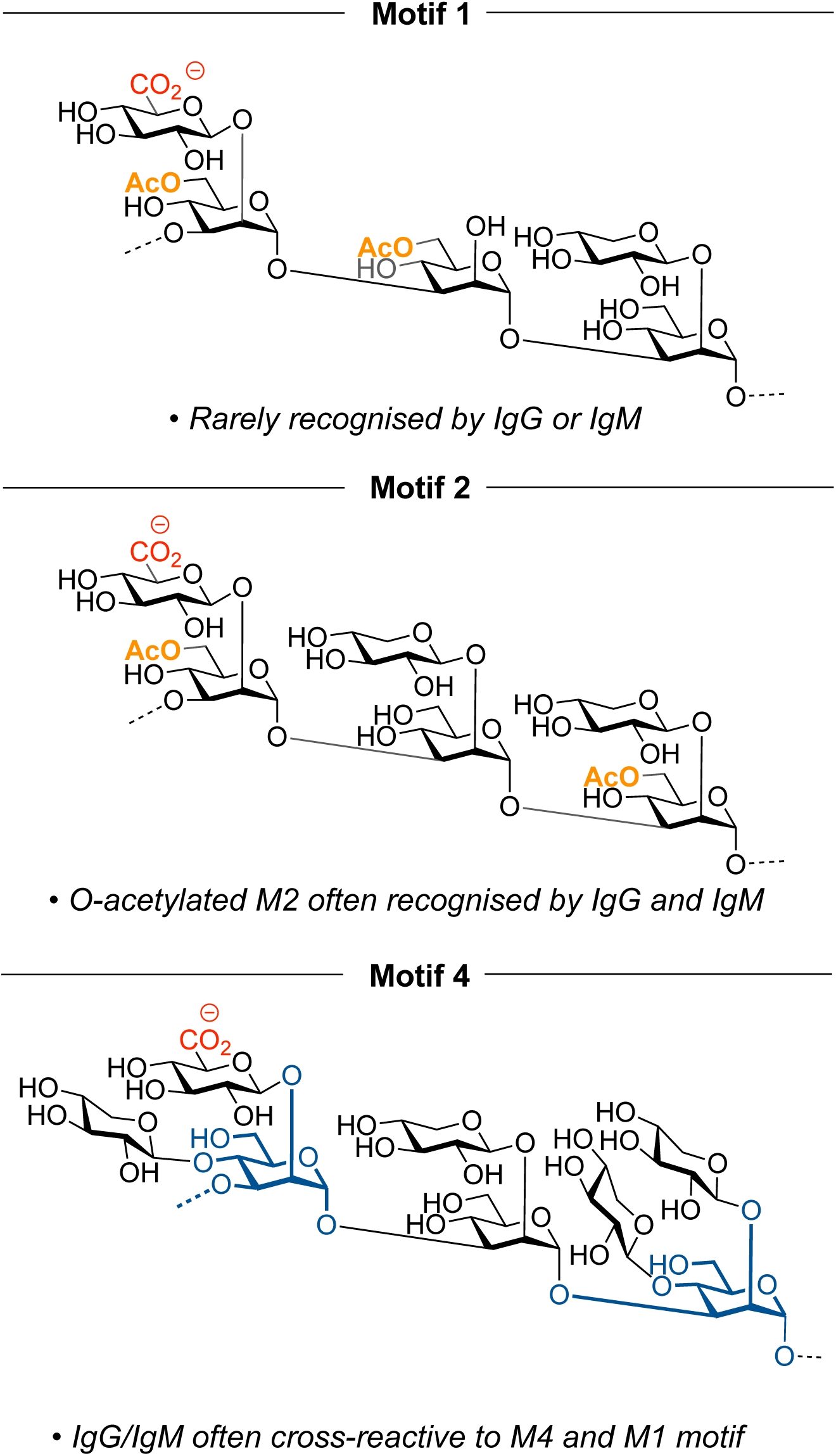
GXM motifs. GXM motifs (M1, M2 and M4), demonstrating the antigenic diversity found in the capsule of *C. neoformans*.

**SI Figure 2.**
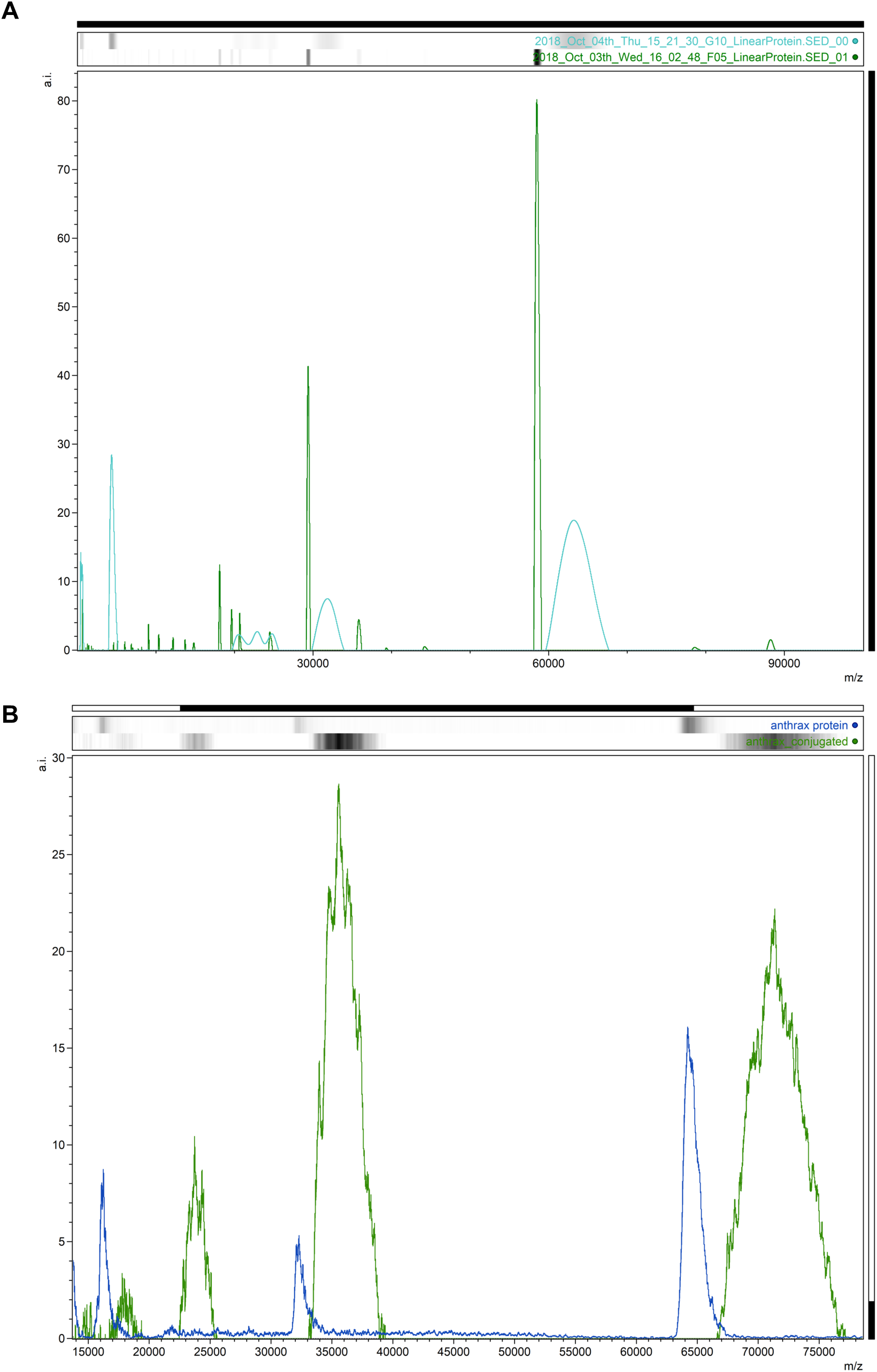
MALDI-TOF of glycoconjugates DECA-CRM197 and DECA-PA63. **a.** DECA-CRM197, shown in green is CRM197 and in blue DECA-CRM197. **b** DECA-PA63 conjugate, shown in green is DECA-PA63 and blue is PA63.

**SI Figure 3.**
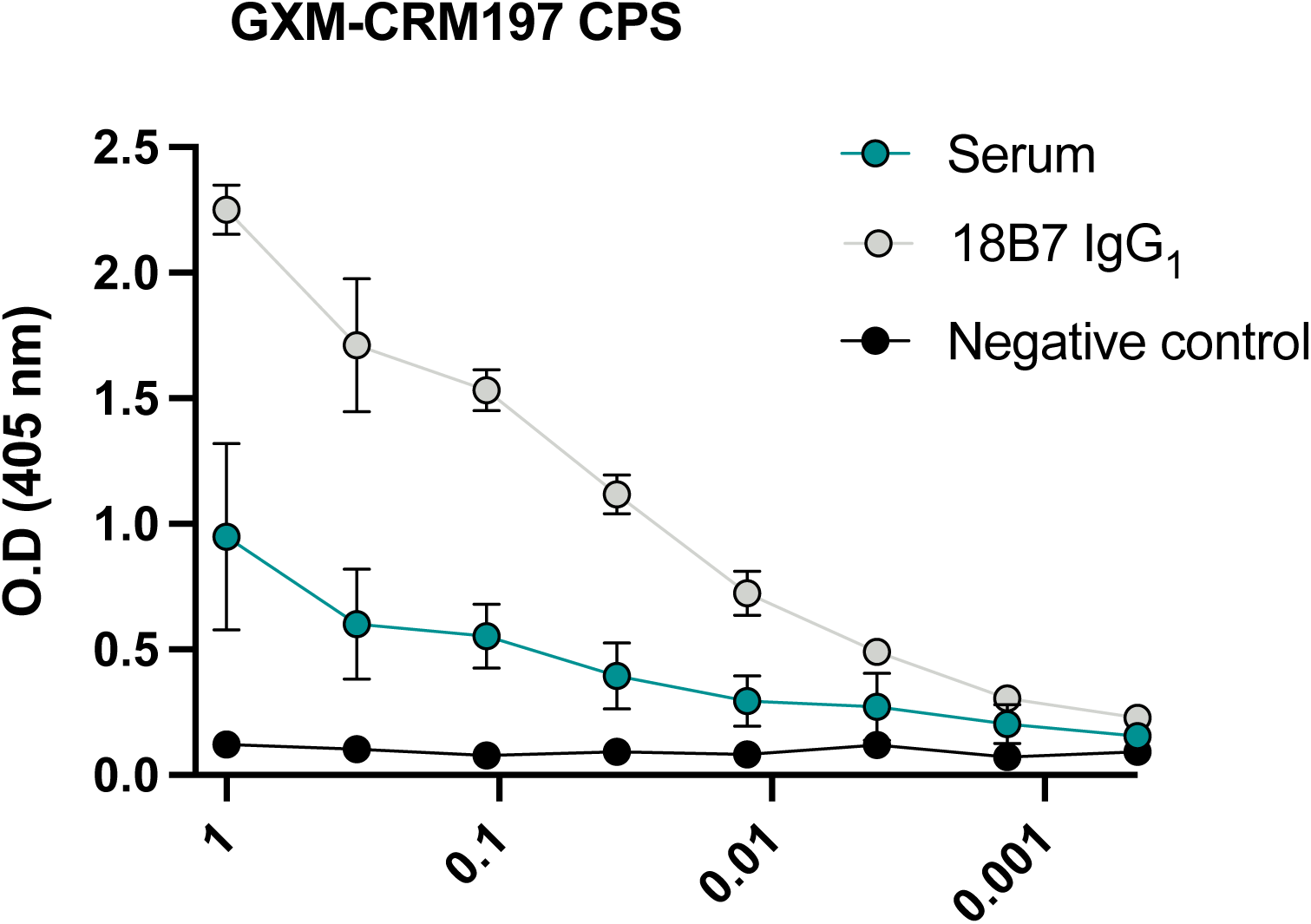
ELISA with capsular polysaccharide and DECA-CRM197 conjugates. Serum from DECA-CRM197 was examined for binding to CPS-coated ELISA plates and exhibited a higher affinity. Positive control: mAb 18B7; Negative control: irrelevant mAb.

**SI Figure 4.**
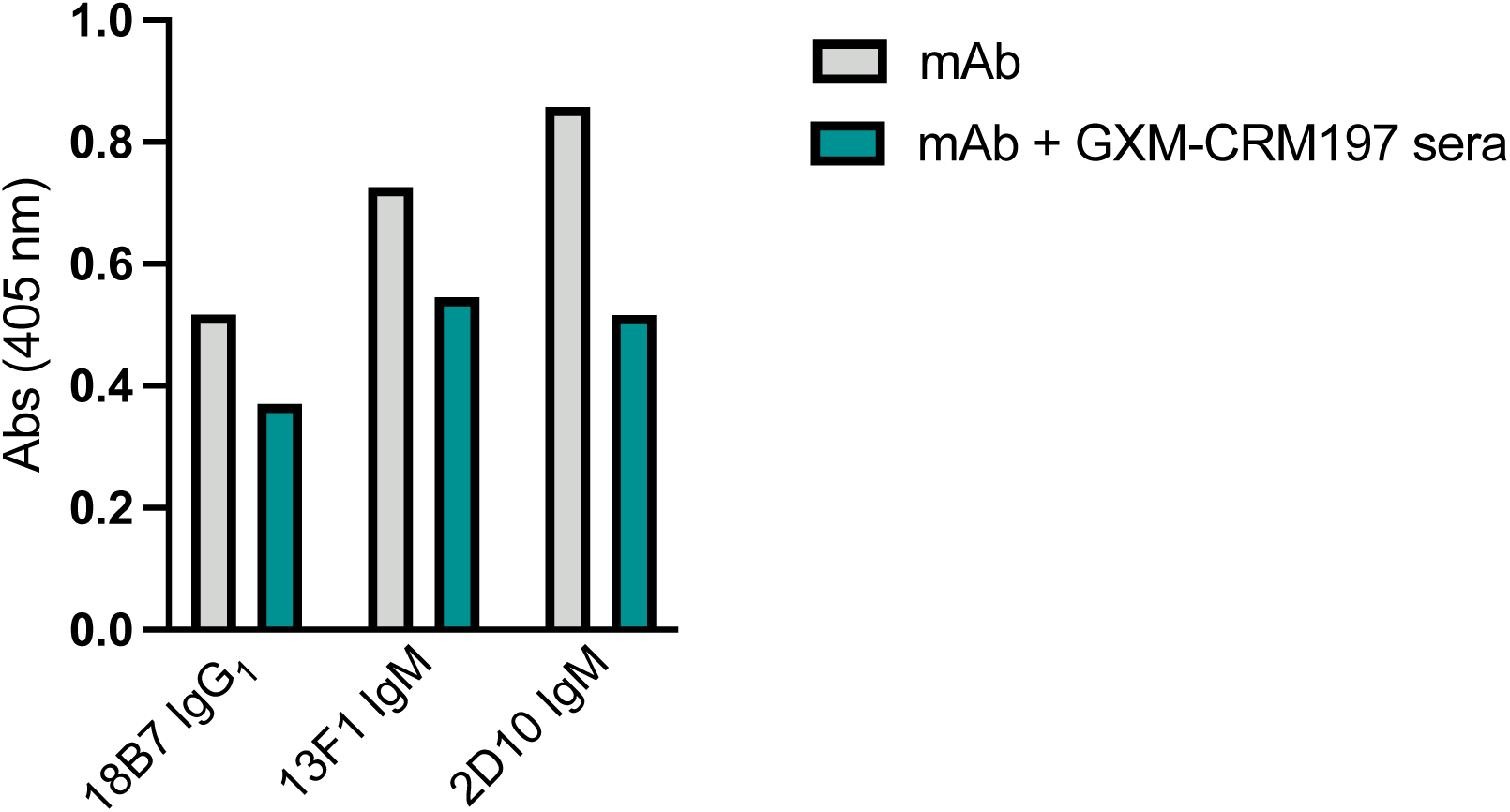
Competition ELISA between mAbs and sera. Serum from DECA-CRM197 mice vs mAbs 18B7 (IgG1), 13F1 (IgM) and 2D10 (IgM) showed that all three mAbs could compete for the same epitopes.

**SI Figure 5.**
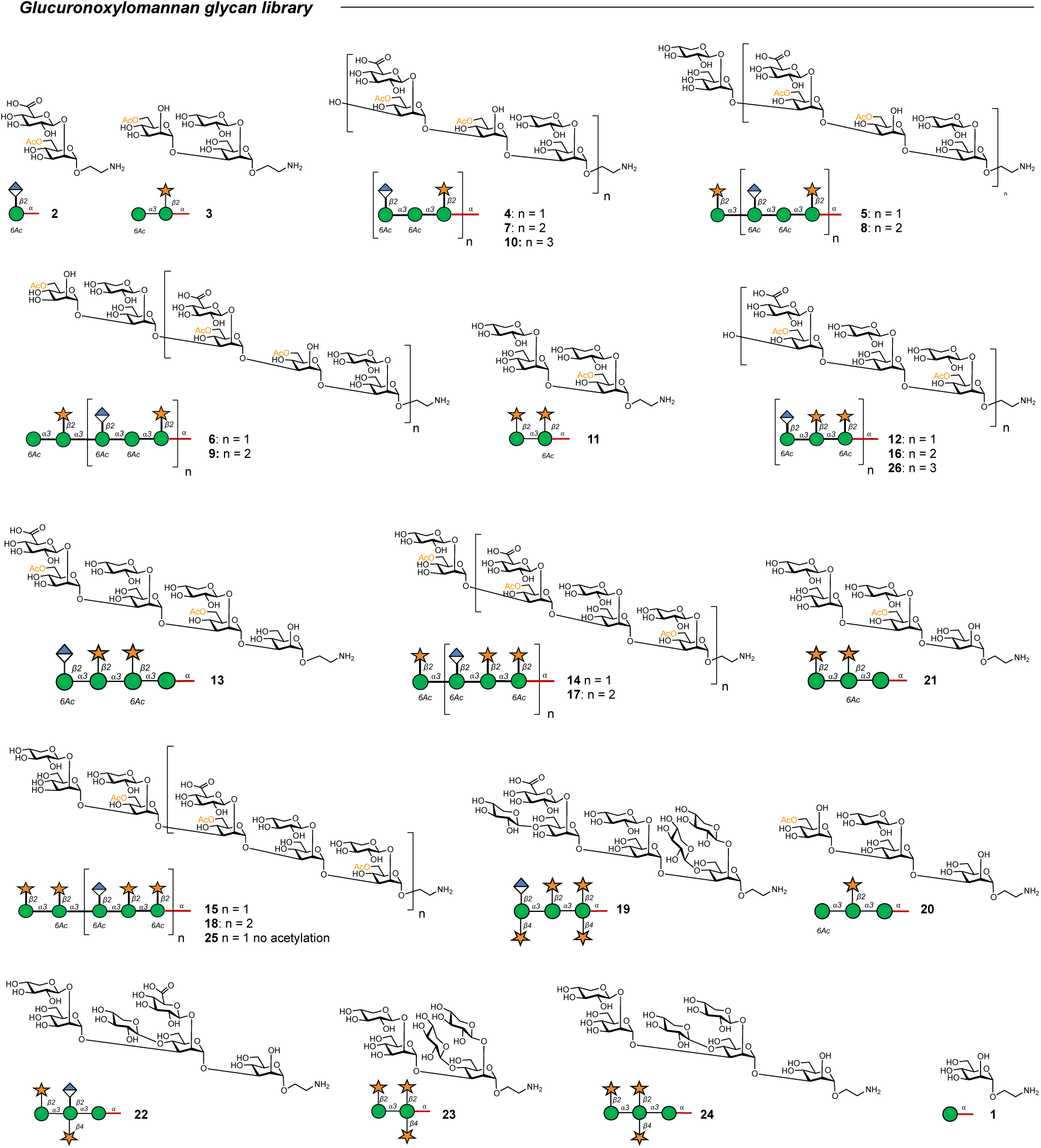
Glucuronoxylomannan glycan microarray library. Full library of the oligosaccharides GXM structures.

